# Multiple pathways to red carotenoid coloration: House finches (*Haemorhous mexicanus*) do not use CYP2J19 to produce red plumage

**DOI:** 10.1101/2024.11.26.625526

**Authors:** Rebecca E. Koch, Christy N. Truong, Hannah R. Reeb, Brooke H. Joski, Geoffrey E. Hill, Yufeng Zhang, Matthew B. Toomey

## Abstract

The carotenoid-based colors of birds are a celebrated example of biological diversity and an important system for the study of evolution. Recently, a two-step mechanism, with the enzymes cytochrome P450 2J19 (CYP2J19) and 3-hydroxybutyrate dehydrogenase 1-like (BDH1L), was described for the biosynthesis of red ketocarotenoids from yellow dietary carotenoids in the retina and plumage of birds. A common assumption has been that all birds with ketocarotenoid-based plumage coloration used this CYP2J19/BDH1L mechanism to produce red feathers. We tested this assumption in house finches (*Haemorhous mexicanus*) by examining the catalytic function of the house finch homologs of these enzymes and tracking their expression in molting birds. We found that CYP2J19 and BDH1L did not catalyze the production of 3-hydroxy-echinenone (3-OH-echinenone), the primary red plumage pigment of house finches, when provided with common dietary carotenoid substrates. Moreover, gene expression analyses revealed little to no expression of *CYP2J19* in liver tissue or growing feather follicles, the putative sites of pigment metabolism in molting house finches. Finally, although the hepatic mitochondria of house finches have high concentrations of 3-OH-echinenone, observations using fluorescent markers suggest that both CYP2J19 and BDH1L localize to the endomembrane system rather than the mitochondria. We propose that house finches and other birds that deposit 3-OH-echinenone as their primary red plumage pigment use an alternative enzymatic pathway to produce their characteristic red ketocarotenoid-based coloration.

## Introduction

Carotenoid-based coloration has been a key area of focus for behavioral and evolutionary biologists for decades, inspiring foundational hypotheses for reliable sexual signaling and for the interplay between physiological processes and display trait expression (Endler 1983; Kodric-Brown and Brown 1984; Hill 1991). The discovery of genes mediating carotenoid-based coloration has recently provided a new approach to testing such hypotheses and understanding how variation in ornament expression may be linked to individual quality (Toews et al. 2017; Hill 2022). Among the best-known of these genes is *CYP2J19*, which encodes a cytochrome P450 enzyme that is involved in the oxidation of yellow dietary carotenoids into red carotenoids that have end rings substituted with at least one ketone group in the C-4 and/or 4’ position (“ketocarotenoids”). Since its first description in canaries (*Serinus canaria*; Lopes et al. 2016) and zebra finches (*Taeniopygia guttata*; Mundy et al. 2016), *CYP2J19* expression has been linked to red carotenoid-based coloration in a diversity of birds (Alonso-Alvarez et al. 2022), such as the long-tailed finch (*Poephila acuticauda*; Hooper et al. 2019), the red-fronted tinkerbird (*Pogoniulus pusillus*; Kirschel et al. 2020), and the red-billed quelea (*Quelea quelea*; Twyman et al. 2018), indicating that *CYP2J19*-mediated red coloration is widespread across avian taxa.

Subsequent functional analyses have revealed that CYP2J19 is necessary, but not sufficient, to catalyze the oxidation of yellow dietary carotenoids to red ketocarotenoids. Gene expression studies of the avian retina, where red ketocarotenoids accumulate in cone oil droplets, identified a second key enzyme: 3-hydroxybutyrate dehydrogenase 1-like (BDH1L; Toomey et al. 2022a). Assays of enzyme function in cell culture revealed that BDH1L is necessary to catalyze the oxidation of the products of CYP2J19 to generate red ketocarotenoids from yellow dietary precursor carotenoids (Toomey et al. 2022a; Figure 1). In the absence of CYP2J19, BDH1L can oxidize dietary carotenoids into yellow canary xanthophylls, pigments that are often observed in avian plumage (Toomey et al. 2022a; Figures 1, 2). The same study also reported a third carotenoid-related protein, TTC39B, that enhances the metabolic conversion of yellow carotenoids to red ketocarotenoids in cell culture (Toomey et al. 2022a). *TTC39B* was discovered through comparison of red and orange morphs of domesticated red-throated parrotfinch (*Erythrura cyaneovirens*), and is also implicated in red ketocarotenoid coloration variation in wild birds (Hooper et al. 2019). These new discoveries present an opportunity to resolve the mechanistic underpinnings of carotenoid-colored trait variation and make inferences about the information content of these signals.

**FIG. 1.**
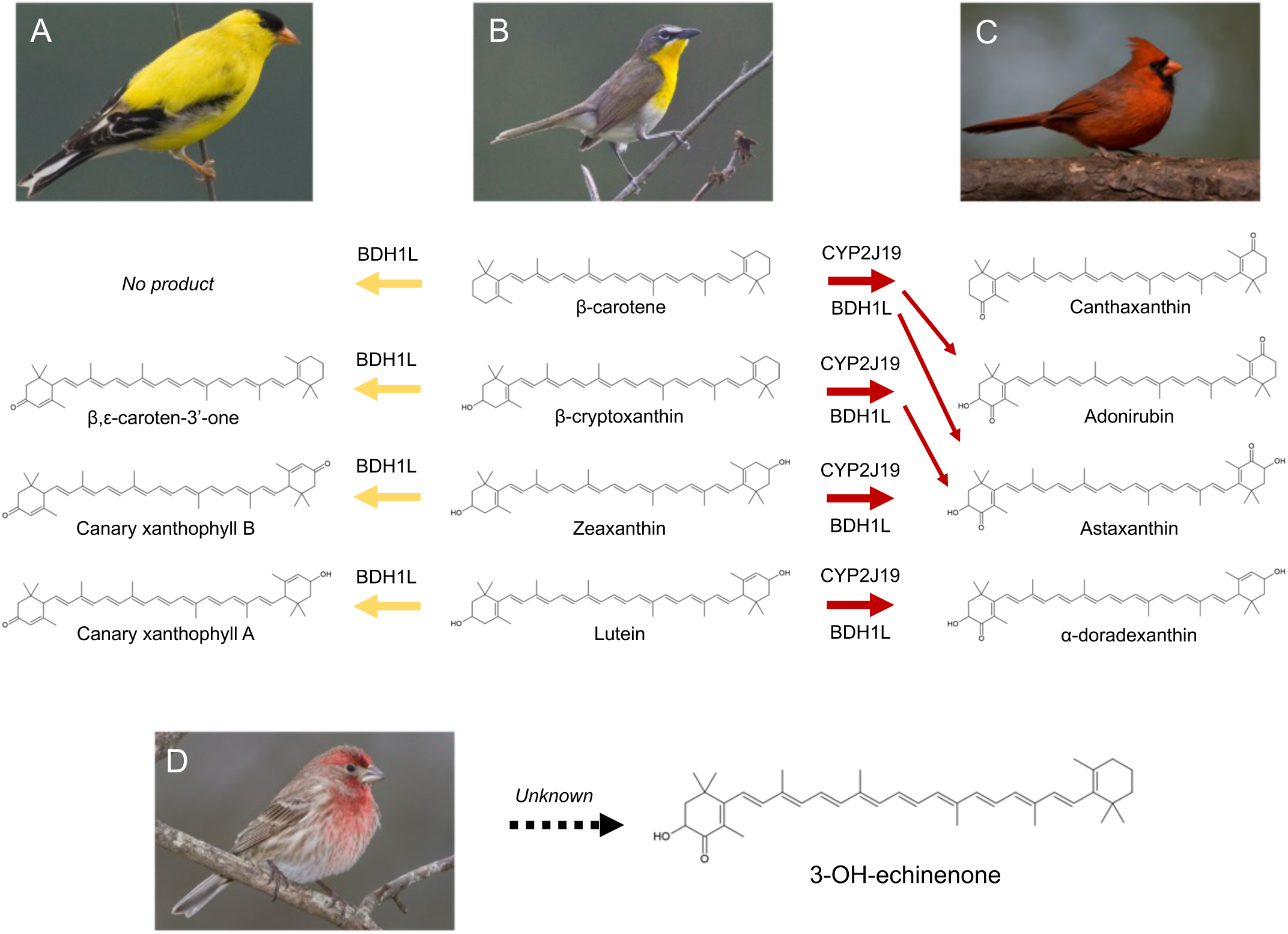
The two best - known carotenoid - metabolizing enzymes in birds, CYP2J19 and BDH1L, can produce a variety of modified yellow carotenoids (left) and red ketocarotenoids (right) when provided different dietary yellow carotenoids (center). Enzymatic activity assays in cell culture testing the house finch homologs of these enzymes found that BDH1L alone catalyzes the conversion of dietary yellow into modified yellow carotenoids, such as canary xanthophylls, which can be found in the plumage of birds like the American goldfinch ( *Spinus tristis*;A). When present with CYP2J19, the two enzymes catalyze the production of red ketocarotenoids from yellow dietary precursors in a stepwise fashion (see Toomey et al. 2022). The four red ketocarotenoids that are the main products of BDH1L and CYP2J19 with the four main dietary carotenoids are found in the red feathers of many birds, including the northern cardinal ( *Cardinalis cardinalis*;C). Some birds do not appear to metabolize carotenoids at all and instead deposit dietary carotenoids directly into plumage ( e.g. yellow - breasted chat, *Icteria virens*;B). However, no combination of these enzymes and carotenoid substrates appears to catalyze the production of the main red ketocarotenoid found in the plumage of male house finches ( *Haemorhous mexicanus*;D). The metabolic pathway(s) birds use to produce ketocarotenoids like 3 - OH - echinenone remain unknown.

House finches (*Haemorhous mexicanus*) are among the best-studied systems for carotenoid-based coloration in wild birds. Male house finches have carotenoid coloration on their head, breast, and rump feathers that varies from dull yellow to orange to brilliant red (Hill 2002). Decades of study suggest that this coloration is an honest signal of male quality and a target of female mate choice (Hill 2002; Toomey and McGraw 2012; Giraudeau et al. 2018). The redness of male feathers is determined by the quantities of red ketocarotenoids, primarily 3-hydroxy-echinenone (3-OH-echinenone), that accumulate in these tissues (Inouye et al. 2001; McGraw et al. 2006). Therefore, the information communicated by house finch plumage coloration is shaped by the specific physiology of carotenoid uptake and ketocarotenoid metabolism from dietary yellow carotenoids, which makes it a useful system in which to explore role of the enzymes that enable the conversion of yellow to red pigments in creating an association between plumage color and individual quality.

While genetic discoveries have provided new tools for studying pigment metabolism, advances in our understanding of the subcellular systems in which these gene products may act has further shaped our perspective on why ornamental coloration varies (Hill 2022; Toomey et al. 2022a). Current theory proposes organismal performance and color expression can be linked through shared dependence on fundamental cellular processes (Hill 2011; Powers and Hill 2021). Early predictions suggested that a main carotenoid-converting enzyme might act within the inner mitochondrial membrane (Johnson and Hill 2013), and CYP2J19 could plausibly fill this role (Hill et al. 2019), thereby linking subcellular processes to ornamental coloration. In support of this hypothesis, the main house finch ornamental ketocarotenoid 3-OH-echinenone has been found in high concentrations in the inner mitochondrial membrane fractions of liver tissue collected from molting male house finches (Ge et al. 2015; Hill et al. 2019). Measures of mitochondrial respiration have also been found to correlate to the redness of plumage (Hill et al. 2019) and the concentrations of 3-OH-echinenone in circulation in molting male house finches (Koch et al. 2024). These observations collectively suggest that the enzymatic conversion of yellow dietary carotenoids to red ketocarotenoids may occur in mitochondria in this species, potentially through the activity of CYP2J19 (Johnson and Hill 2013; Hill et al. 2019).

Despite the house finch being considered a model for testing the CYP2J19 pathway (now known to also involve BDH1L) for metabolizing yellow dietary pigments to red ornamental pigments (Hill et al. 2019), no studies have established CYP2J19/BDH1L expression or function in relation to the production of red feather pigments in house finches. Here, we test three key predictions regarding the role of the CYP2J19/BDH1L pathway in the production of red ketocarotenoid pigments in the house finch: 1. *CYP2J19* and *BDH1L* are expressed in male house finches at the putative sites of plumage pigment metabolism; 2. CYP2J19 and BDH1L catalyze the formation of 3-OH-echinenone, the major ketocarotenoid pigmenting red house finch feathers; and, 3. CYP2J19 localizes to and functions within the mitochondria. Our studies focus on the house finch, but our tests hold implications for the other bird species that use 3-OH-echinenone or other ketocarotenoids with similar structures as their primary red plumage pigment.

We tested these predictions using a combination of gene expression analyses, enzymatic activity assays, and protein localization studies. We first examined the expression patterns of *CYP2J19* and *BDH1L* in retinas, livers, and growing feather follicles of molting and non-molting wild male house finches by RNA sequencing. We then cloned the house finch homologs of all three genes and performed enzymatic activity assays in cell culture to assess the products formed by these enzymes when provided with different carotenoid substrates. We also created versions of these cloned genes tagged with fluorescent markers that, in combination with markers localizing specifically to different subcellular locations, allowed us to visualize where in the cell each gene product localized. While we focused our analyses on house finches, our findings have important implications for understanding the evolution of red carotenoid-based coloration in Aves, and we place our results from the house finch within the broader context of carotenoid metabolism across the phylogeny of birds.

## Materials and Methods

### Tissue Sampling

We captured house finches in traps at established feeding sites in Auburn, Alabama, USA, according to the methods described in Hill (2002). To acquire feather follicles and other tissues from house finches that were actively growing feathers, in August 2021 we captured male finches, euthanized the birds, and collected blood samples (immediately centrifuged to separately store plasma from other blood components), whole liver, skin containing growing follicles, and whole eye samples. We immediately froze the samples in liquid nitrogen and then stored them at -80°C until extraction and analysis. We also plucked fully grown, colored feathers from the breast and rump of adult males and stored these in the dark at ambient temperature prior to pigment analysis. To acquire growing feather follicles from non-molting birds, we captured birds on January 11 and 12, 2022, determined sex using plumage characteristics, plucked a region of red breast feathers to stimulate feather regrowth, and held the birds in large outdoor aviaries as they regrew plucked feathers (see Koch et al. 2024 for details of the outdoor aviaries). Eleven days after we plucked birds, these finches were euthanized and growing feather follicles were collected from each bird and frozen. We also captured two male purple finches (*Haemorhous purpureus)*, the sister species to the house finch, in Auburn, Alabama on March 15, 2023 using the same methodology, and plucked 15 red breast and rump feathers from each bird for carotenoid analyses. All procedures involving live animals were approved by the Auburn University Institutional Animal Care and Use Committee with state of Alabama and United States collecting permits.

### RNA extraction, Sequencing, and Processing

We extracted RNA from portions of frozen follicle, liver, and retina tissues using the TRIzol Reagent (Thermo Fisher Scientific Inc., Waltham, MA, USA). We first dissected out the relevant portions of each tissue: the retina from one eye, 250-500 mg of liver tissue, and 7-20 follicles from each skin sample. In an effort to sample follicles at the same developmental stage, we limited our sample to follicles that were 7-8 mm in length. We homogenized each sample in 1 mL TRIzol Reagent with 0.1 g of zirconia beads (ZROB10; Next Advance, Inc., Troy, NY, USA) in a Beadbug homogenizer (Benchmark Science, Inc., Sayreville, NJ, USA) for 180s at 4 kHz. We then proceeded with RNA extraction according to the manufacturer’s guidelines, with the addition of 1 μL glycogen (Thermo Fisher Scientific Inc.). After this initial RNA extraction, we removed residual DNA from each 43.5 μL RNA sample by adding 1.5 μL Turbo DNase and 5 μL Turbo DNase Buffer (Invitrogen TURBO DNA-free Kit, Thermo Fisher Scientific Inc.), then incubating for 30 mins at 37C. We then re-extracted RNA from this DNAse-treated sample: first, we added 150 μL molecular grade water and 200 μL chloroform, mixed by vortex, centrifuged, then collected the aqueous fraction to a new tube. Next, we added 17.5 μL sodium acetate (pH 5.2 3M; Alfa Aesar, Ward Hill, MA, USA), 1 μL glycogen, and 600 μL ethanol, and incubated the samples for 20 mins at -20C. We centrifuged the samples and removed the supernatant, and finally washed the pellets twice with 80% ethanol before air-drying and resuspending in 25 μL molecular grade water. RNA samples were stored at -80C until further analysis. For sequencing, we submitted 100 ng of total RNA to the Clinical Genomes Laboratory at the Oklahoma Medical Research Foundation (OMRF; Oklahoma City, Oklahoma, USA). OMRF prepared mRNA sequencing libraries using the xGen RNA Lib Prep Kit (Integrated DNA Technologies, Coralville, IA, USA) with the NEB poly-A selection kit (New England Biolabs, Ipswich, MA, USA) and sequenced the mRNA libraries as 150 bp paired-end reads on an Illumina NovaSeq 6000. In total, we obtained expression data from three retina, three liver, and four follicle samples from molting males, and three follicle and three liver samples from non-molting males.

We received demultiplexed sequencing reads in FASTQ format, evaluated quality with FastQC (v. 0.11.5; Andrews 2010), and trimmed the paired reads using Trim Galore, set to trim adaptors and low quality bases (Phred score <5) and discard reads shorter than 36 bp (v. 0.6.0; Krueger 2021). We used Hisat2 (v. 2.1.0; Kim et al. 2019) to align the reads to a house finch genome (GenBank: GCA_027477595.1), and sorted and indexed the alignments using SAMtools (v. 1.11-GCC-10.2.0; Danecek et al. 2021).

We quantified gene expression from the alignments first by using the featureCounts function within the Rsubread package (v. 2.2.3; Liao et al. 2019) in R (v. 4.2.1; R Core Team 2023) to assign raw read counts to annotated genes. We then used DESeq2 (v. 1.40.2; Love et al. 2014) to evaluate significantly differentially expressed genes, excluding genes with fewer than 10 total counts. We performed this analysis separately for data from liver and from follicles; for each tissue type, we specified season (molt or non-molt) as a variable in the design formula, with non-molt as the reference group. We also transformed read counts for each gene within a tissue using the size factor and rlog normalization tools within DESeq2 to obtain expression values that account for differences in sequencing depth and heteroskedasticity (Tables S1, S2).

Lastly, we visualized expression of focal genes by importing the house finch genome and annotation file along with the aligned reads for each sample into Integrative Genomics Viewer (IGV; v. 2.12.3; Robinson et al. 2011). Neither *CYP2J19* nor *BDH1L* are specifically annotated in the current house finch genome, however a BLAST (Camacho et al. 2009) search with the *Serinus canaria* homolog sequences (NCBI Reference Sequences XM_050977223 for *CYP2J19* and XM_018918048 for *BDH1L*) revealed that the genes map to two “uncharacterized loci”: LOC132330886 (*CYP2J19*) and LOC132324410 (*BDH1L*) in the current house finch annotation (GCF_027477595.1-RS_2023_09).

### Gene Cloning

For *CYP2J19*, we assembled a full transcript of the house finch coding sequence from two partial PCR products amplified from retinal cDNA, using Gibson assembly to simultaneously insert the transcripts into the first position of a bicistronic expression construct that encoded either a green or red fluorescent protein in the second position (pCAG-[first position]-2A-GFP or pCAG-[first position]-2A-dsRed, respectively). We could not successfully amplify the entirety house finch *BDH1L* via PCR (likely because of high GC content), but instead used Gibson assembly to build the assemble the predicted coding sequence using codon-optimized synthesized fragment (Integrated DNA Technologies, Inc., Coralville, IA, USA) of the 5’ position of the transcript and 3’ fragment amplified by PCR from follicle cDNA. Lastly, were able to amplify the predicted full-length transcript of the house finch *TTC39B* coding using PCR on follicle cDNA and ligate this into a third expression construct. We verified the sequences of all three constructs using Oxford Nanopore long read sequencing of whole plasmids via plasmidsaurus (SNPsaurus LLC, Eugene, OR, USA). The sequences of primers and synthesized DNA fragments used in this study are listed in Table S3.

### Enzymatic Activity Assays in Cell Culture

We performed cell-based assays of enzymatic activity by transiently transfecting cultured HEK-293 cells (ATTC, CRL-1573), grown to 70% confluency in a Dulbecco’s Modified Eagle Medium mixture (cytiva HyClone DMEM, Wilmington, DE, USA; 10% Fetal Bovine Serum, Fisher Scientific; 100 U/mL penicillin and streptomycin, Gibco; and 1x GlutaMAX, Gibco, Thermo Fisher Scientific), with one or more constructs using polyethylenimine (PEI, Polysciences, Inc.; Warrington, PA, USA, 23966-2).

Briefly, for transfection, we first added serum-free DMEM to a sterile tube in a volume equaling 10% of the total volume of media provided to the cells to be transfected; we then added plasmid DNA at a quantity of 3 μg per 0.5 x 10^6^ cells, and mixed in PEI at a ratio of 3 μg PEI : 1 μg DNA and incubated 15 mins at room temperature before adding dropwise to cultured cells. For each assay, we also transfected control cells with a construct containing fluorescent proteins in both positions (pCAG-GFP-2A-dsRed). We incubated transfected cells for 48 hours and verified construct expression by visualizing the expression of the fluorescent proteins.

In each assay, we provided the transfected cells with media supplemented with specific carotenoid pigments—purified β-cryptoxanthin, purified zeaxanthin, purified lutein, or β-carotene—to serve as substrates for heterologously expressed enzymes. We ensured that enzymes were provided with different substrates that together presented the three different configurations of end rings found in the main dietary carotenoids: an unsubstituted β-ring (β-cryptoxanthin, β-carotene), a β-ring hydroxylated at the C3 position (β-cryptoxanthin, zeaxanthin), and an ε-ring (lutein). We extracted zeaxanthin, lutein, and β-carotene from carotenoid beadlet samples provided by DSM (dsm-firmenech, Stroe, Netherlands; OPTISHARP [5% zeaxanthin - 5003563004]; β-carotene 10% [0489999004]; FloraGLO [10% lutein - 5011868022]), and β-cryptoxanthin from freshly squeezed mandarin juice. Carotenoids extracted from the mandarin juice were also saponified for 6 hours with 0.2M NaOH in methanol and re-extracted before further processing. From these extractions, we purified the all-*trans* forms of β-cryptoxanthin, lutein, and zeaxanthin using high performance liquid chromatography (HPLC) separation methods described below. We did not further purify the β-carotene extract because it was found to primarily contain the all-*trans* form of this carotenoid. The purified substrate carotenoids were added individually to the cell culture media at concentrations of 0.8-1.2 μg/mL with polysorbate 40 (Tween 40; AC334142500; Acros Organics, Thermo Fisher Scientific, Inc.) at a concentration of 0.035%. 18-20 hours after the additional of the carotenoid substrates, we collected and pelleted experimental cells via centrifugation, washed the pellet twice with phosphate-buffered saline, and then stored the washed pellet in the dark at -80C until further analysis.

### Carotenoid Extraction and Analysis

To extract carotenoids from the collected cell pellets, we first resuspended each pellet in 500 μL of 0.9% NaCl containing 0.1 g of zirconia beads. We disrupted the cells using the Beadbug homogenizer for 30 s at 4 kHz, then added 250 μL of 100% ethanol, mixed by vortex, added 500 μL of hexane:*tert*-butyl methyl ether (1:1, vol:vol; hexane:MTBE), and ground once more for 30 s at 4 kHz. We centrifuged the cell homogenate at 10,000 g for 3 minutes, then collected the upper solvent layer into a 2 mL glass vial and completely evaporated it under a constant stream of nitrogen. We extracted carotenoids from plasma samples using a similar process: to 10 μL of plasma, we added 250 μL of 100% ethanol followed by 250 μL hexane:MTBE, vortexed, then centrifuged at 10,000 g for 3 minutes. We then transferred the upper solvent layer to a glass vial and dried under nitrogen as described above. To extract carotenoids from feather samples, we trimmed colored barbs into a 1.5 mL screw-cap tube and added 1 mL of 100% methanol along with 0.1 g of zirconia beads. We ground the feather barbs in the Beadbug homogenizer at 4 kHz for 5 minutes, then centrifuged at 10,000 g for 3 minutes and extracted supernatant to a glass vial. We repeated this process two more times to extract additional carotenoids remaining in the sample, then dried the collected extract under a stream of nitrogen gas.

For carotenoid analysis via HPLC, we resuspended each dried extract in 120 µL of mobile phase—acetonitrile:methanol:dichloromethane (44:44:12, vol:vol:vol)— and injected 100 μL into an Agilent 1200 series HPLC with a YMC carotenoid column maintained at 30C (5.0 µm, 4.6 mm × 250 mm, YMC, CT99S05-2546WT). We eluted samples at a constant solvent pump rate of 1.2 mL/min and with a mobile phase of acetonitrile:methanol:dichloromethane (44:44:12) for 11 minutes, which then increased to acetonitrile:methanol:dichloromethane (35:35:30) from 11-21 minutes, and continued as isocratic conditions until 35 minutes. We monitored sample elution using a UV-Vis photodiode array detector at 445 or 480 nm, and we identified carotenoids by comparing to standards of astaxanthin, canthaxanthin, zeaxanthin, lutein, and β-carotene (dsm-firmenech, Stroe, Netherlands) or from published accounts (Inouye et al. 2001; Britton et al. 2004; Potticary et al. 2020).

### Candidate Carotenoid Genes

In addition to testing our three focal coloration genes, we leveraged our gene expression results to explore new candidate genes that may be involved in carotenoid metabolism. In total, we cloned (using methods as described above; Table S3) and tested nine different genes with at least two different carotenoid substrates each (Table S4). These genes included three cytochrome P450 enzymes: *CYP2C19*, *CYP26B1*, and *CYP2J2* (sometimes designated as *CYP2J40*), the latter of which is adjacent to *CYP2J19* on the house finch genome and is broadly expressed across house finch tissues. Additional genes we tested included hydroxysteroid 11-beta dehydrogenase 1 (*HSD11B1*), 3-hydroxybutyrate dehydrogenase 1 (*BDH1*), short chain dehydrogenase/reductase family 42E member 2 (*SDR42E2*), fatty acid 2-hydroxylase (*FA2H*), pyridine nucleotide-disulphide oxidoreductase domain 2 (*PYROXD2*), and retinol dehydrogenase 10 (*RDH10*; Table S4). We also co-expressed these candidate genes along with *BDH1L* to test whether they may act specifically on the modified products of BDH1L with carotenoid substrates. All genes were cloned from house finch sequences except *CYP2C19*, which we cloned from Gouldian finch (*Erythrura gouldiae*) DNA based on a previous analysis of Gouldian finch color morphs. Translated Gouldian finch *CYP2C19* is identical to house finch CYP2C19 at 454/494 (91%) amino acid position, with an estimated amino acid similarity of 95.5% (EMBOSS Needle, European Molecular Biology Laboratory; Madeira et al. 2022).

### Enzyme Localization

To trace the subcellular localization of house finch CYPJ19, BDH1L, and TTC39B, we generated expression constructs for fusions of each protein with the mNeonGreen (Shaner et al. 2013) or mCherry (Shaner et al. 2004) fluorescent proteins. To do this, we amplified coding sequencing of each protein, from the expression constructs described above, by PCR, and subcloned these by Gibson assembly (E2621, New England Biolabs, Inc.) into a pCAG vector with the coding sequence of the fluorescent protein in-frame at the C-terminus of the protein. To label mitochondria and ER we used the mCherry-Mito-7 (a gift from Michael Davidson - Addgene plasmid # 55102;http://n2t.net/addgene:55102;RRID:Addgene_55102) and mCherry-ER-3 (a gift from Michael Davidson - Addgene plasmid # 55041; http://n2t.net/addgene:55041; RRID:Addgene_55041) constructs (Olenych et al. 2007). We cultured HEK293 cells on poly-L-ornithine (A004C, Millipore Sigma Inc., Burlington, MA, USA) coated glass coverslips and transfected these cells with various combinations of these constructs following the culture conditions and transfection protocols described above. 16 to 36 hours after transfection, we removed culture media and fixed the cells with 4% paraformaldehyde in phosphate buffered saline (PBS) for 10 minutes at room temperature. We then washed the cells 3x with, counterstained with DAPI (1 ug/mL in PBS, D95421MG, Sigma Aldrich Inc., Burlington, MA, USA), mounted the coverslips on glass slides with Fluoromount G (0100-01, SouthernBiotech Inc., Birmingham, AL, USA) mounting media, and sealed the edges of the coverslips with translucent nail polish. We imaged the cells at 60x magnification with a Zeiss LSM800 laser confocal microscope and processed images with the ZEN software package (ver. 3.2, Carl Zeiss GmbH).

### Phylogenetic comparison

To explore phylogenetically distribution of metabolized carotenoid types across bird species, we characterized the presence of seven different classes of carotenoid pigments detected in bird feathers: unmodified yellow dietary carotenoids (e.g. β- carotene, zeaxanthin, lutein), modified yellow carotenoids (e.g. canary xanthophylls), symmetric β,β-C4-ketocarotenoids (e.g. astaxanthin, canthaxanthin), asymmetric β,ε-C4-ketocarotenoids (e.g. α-doradexanthin), asymmetric β,β-C4-ketocarotenoids (e.g. 3-OH-echinenone), asymmetric β,Ψ-C4-ketocarotenoids (e.g. 4-oxo-rubixanthin), and retrocarotenoids (e.g. rhodoxanthin; Figure 2). Here, a ketocarotenoid is considered symmetric if it has ketone groups at the same positions on both rings, irrespective of locations of hydroxyl groups; adonirubin, for example, is a symmetric β,β-C4-ketocarotenoid despite a hydroxyl group on only one end ring. We collected information on feather carotenoid content across 230 bird species from two published reviews that tabulated such data (McGraw 2006; Toomey et al. 2022b) and from our own data on purple finch feathers from this study, then mapped reported carotenoid types per species on to the least-squares consensus phylogenetic tree from a set of 100 trees from BirdTree.org (Jetz et al. 2012; Jetz et al. 2014), using the *ggtree* (Yu et al. 2017) and *tidyverse* (Wickham et al. 2019) packages in R (v. 4.3.2) within RStudio (v. 2023.09.1+494; RStudio Team 2023). To focus more closely on finches in the genus Fringillidae, we also generated a second tree using updated phylogenetic relationships as reported in Ligon et al. (2016).

**FIG. 2.**
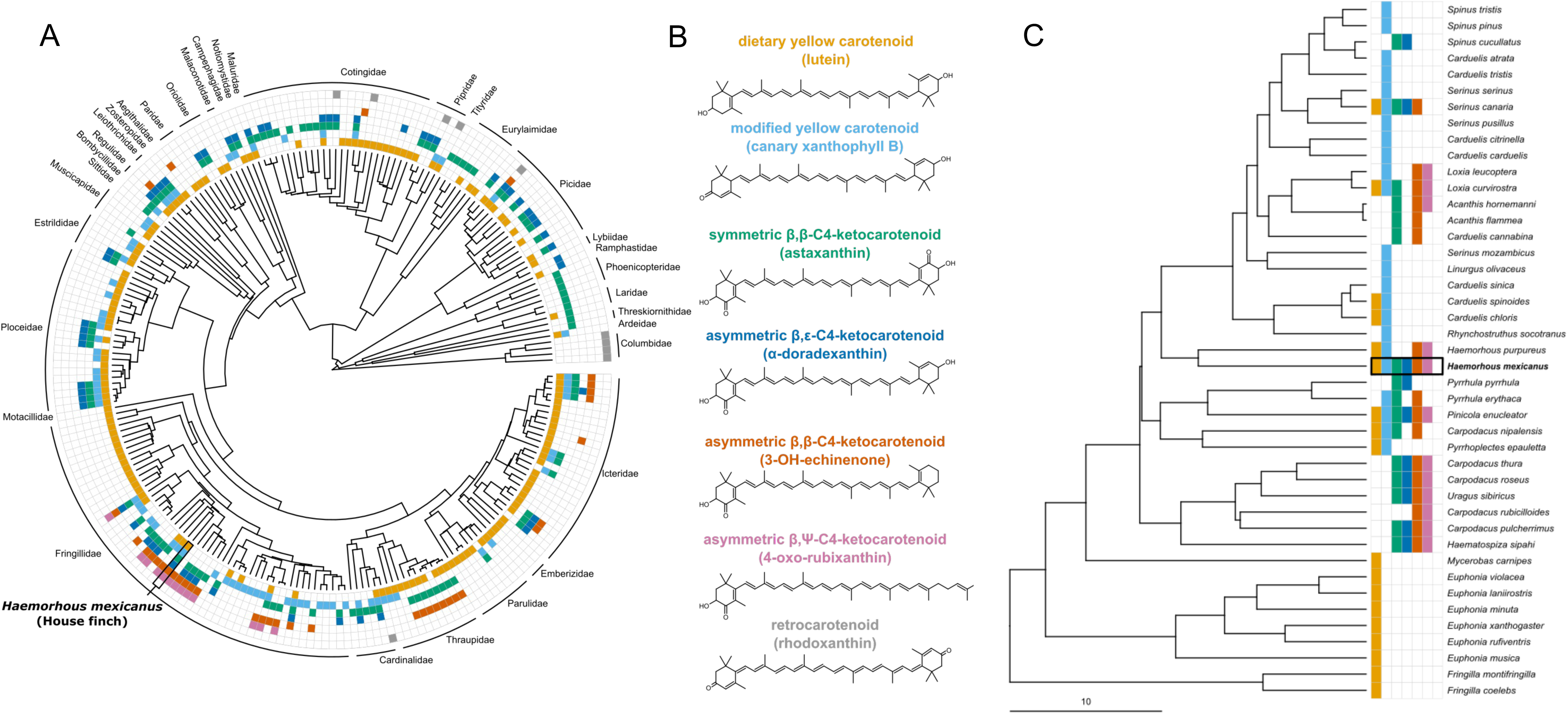
We categorized different carotenoid pigments that have been detected in colored bird plumage into seven main structural class es (B), based on the properties and substitution patterns of their end rings; one example of a carotenoid from each category is shown. We then mapped when carote noi ds of each of these structural categories has been reported in species across a phylogeny of avian taxa (A), with special focus on Fringillid finches (C). Note that negati ve data does not necessarily mean the absence of that type of carotenoid in a taxon, only that it was not reported in the literature we examined.

## Results

### Gene Expression

Across all samples, RNA reads had an overall alignment rate to the house finch genome of 61.7% (± 3.7%; mean ± standard error). *BDH1L*, *TTC39B*, and *CYP2J19* all showed expression in male house finch retina and growing follicle tissue, with little to no expression in liver tissue (Figures 3, Figures S1-2). However, when we investigated the mapping of reads to the genome, we found that the majority of reads from liver or follicle tissue that were assigned to *CYP2J19* map to the gene’s 3’ UTR, and reads were essentially absent from the exons containing the protein-coding sequence of *CYP2J19* (Figure 3). This finding is consistent with our observation that we could successfully amplify full-length house finch *CYP2J19* transcripts via PCR from retinal cDNA, but not follicle or liver cDNA; if only the 3’ UTR of *CYP2J19* is expressed in the liver and growing follicles, cDNA from these tissues would not contain the protein-coding region we targeted. Retinal expression of *CYP2J19* is expected due to its function in metabolizing red carotenoids to pigment the oil droplets of the red-sensitive cone photoreceptor (Toomey et al. 2022a).

**FIG. 3.**
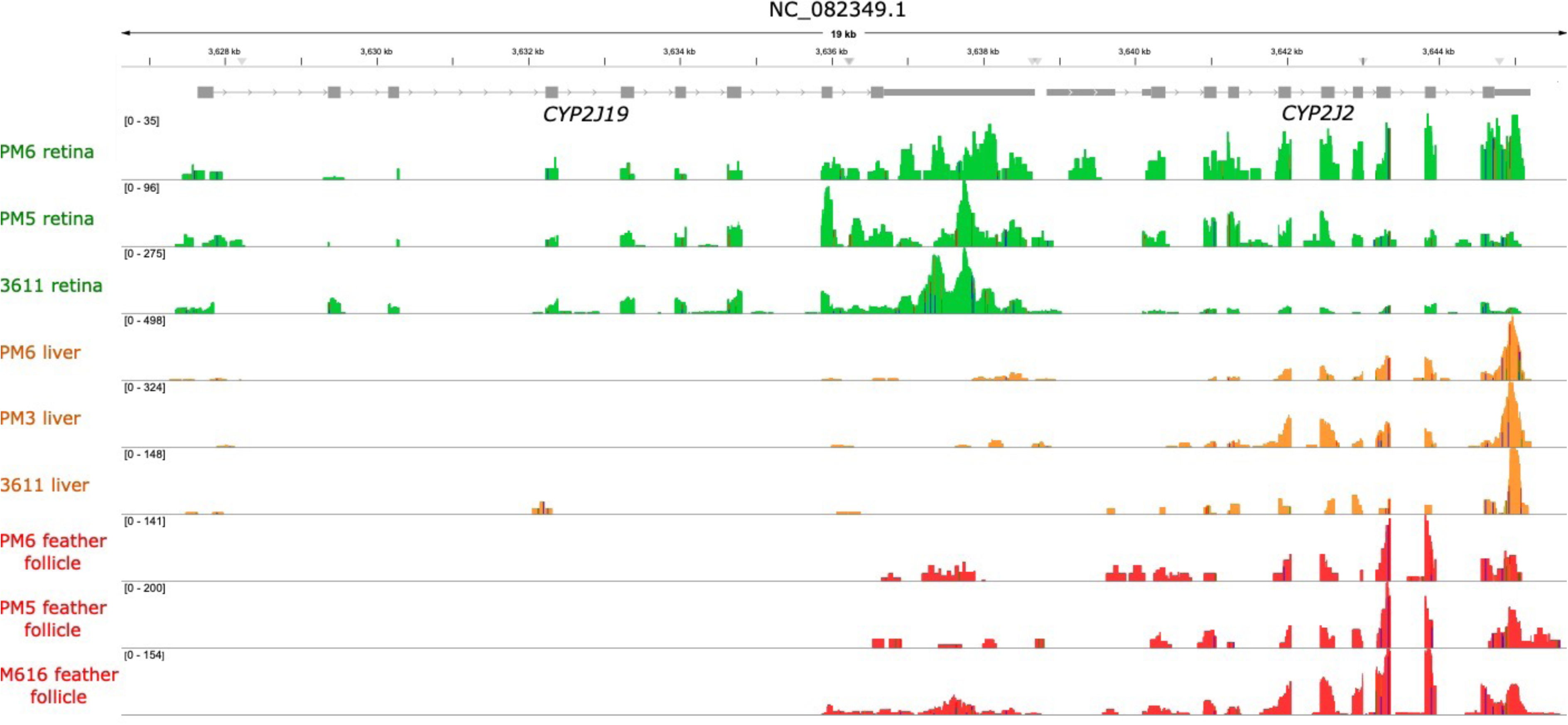
Mapping of RNA - seq reads to the *CYP2J19* locus of Chromosome 8, from male house finch retina (top, green), liver (middle, orange), and growing feather follicle (red, bottom) samples. The exons of *CYP2J19* are designated in grey boxes. As expected due to its involvement in avian vision, *CYP2J19* appears expressed across its length in retinal samples. However, in both the liver and the follicle, some reads align to the 3’ UTR of the gene, but the p rot ein - coding region of *CYP2J19* does not appear expressed.

Differential gene expression analyses identified 1165 follicle and 25 liver genes significantly enriched in molting males compared to non-molting males (Tables S1, S2). However, the expression of *CYP2J19, BDH1L,* and *TTC39B* did not significantly differ between molting and non-molting samples for either tissue.

### Enzymatic Activity Assays in Cell Culture

While the follicle and liver tend to be considered the most likely locations of ketocarotenoid metabolism in birds, our gene expression observations do not preclude the possibility that house finches metabolize carotenoids in a different tissue. As such, we assayed the enzymatic activity of house finch CYP2J19 and BDH1L to determine if these enzymes can catalyze the production of 3-OH-echinenone, the major ketocarotenoid pigmenting red house finch plumage. We found that house finch CYP2J19, BDH1L, and TTC39B exhibited the same properties as previously described for these enzymes from chickens (*Gallus gallus*; Toomey et al. 2022a). For example, when we provided a zeaxanthin substrate to cells co-transfected with house finch *CYP2J19* and *BDHL1,* they produced the ketocarotenoid astaxanthin (Figure 4), while cells transfected with *CYP2J19* alone catalyzed the formation of an oxidized yellow carotenoid, and cells expressing *BDH1L* alone primarily catalyzed the formation of a product consistent with the modified yellow carotenoid canary xanthophyll B (Figure S3). The products of house finch CYP2J19 and BDH1L with lutein and β-carotene substrates, α-doradexanthin and canthaxanthin, respectively, were also identical with those previously reported for the chicken homologs of these enzymes (Figure 4; Toomey et al. 2022a). House finch TTC39B also appears to enhance ketocarotenoid metabolism, as has been reported for chickens (Toomey et al. 2022a). Co-transfection of house finch *TTC39B* with house finch *CYP2J19* increases the relative amount of CYP2J19 product produced (Figure S4); as such, we co-transfected *TTC39B* with our other focal and candidate genes to enhance our ability to detect and visualize new products.

**FIG. 4.**
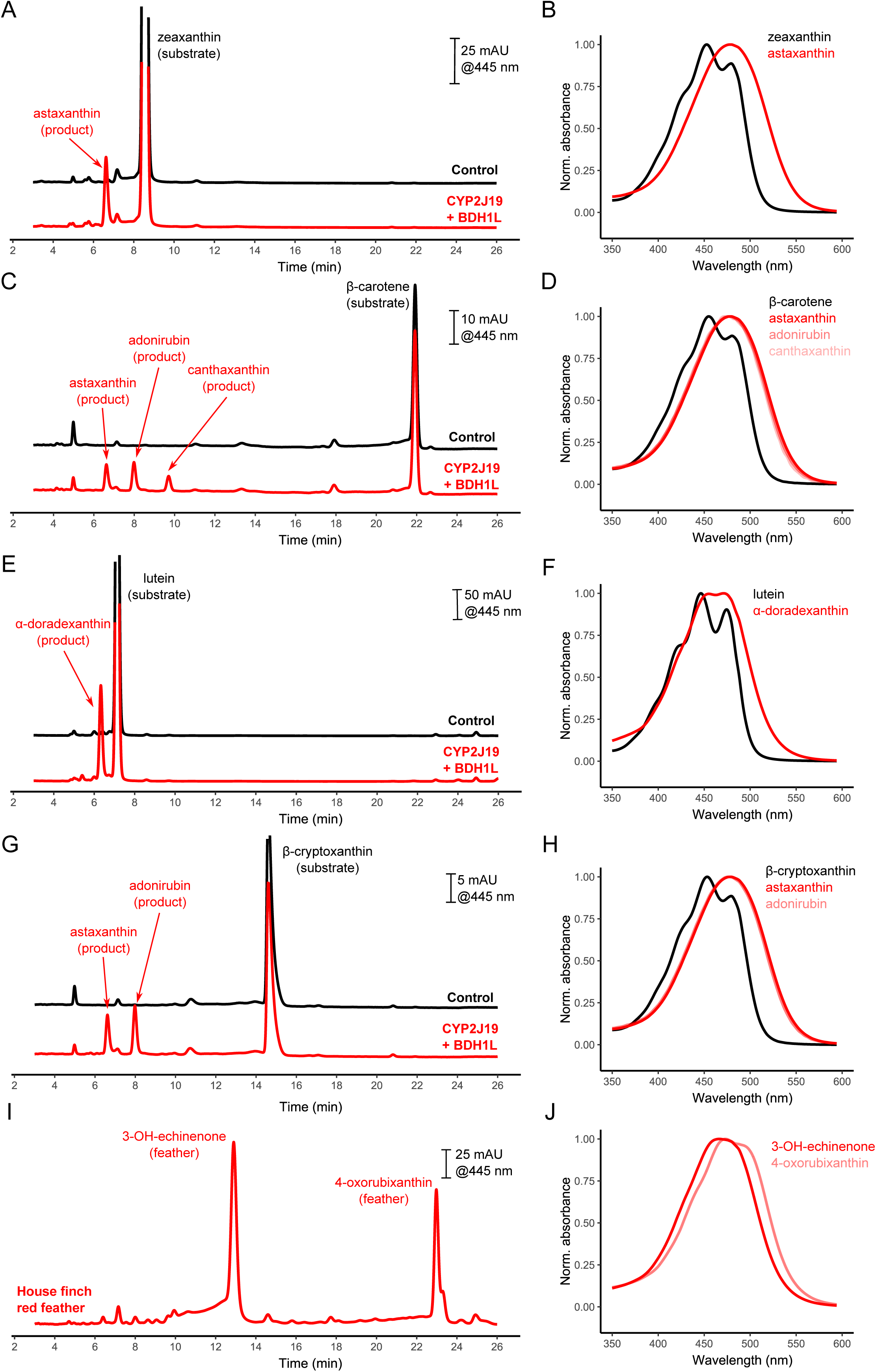
We cloned the house finch homologs of three carotenoid - related genes, *CYP2J19, BDH1L,* and *TTC39B*, to validate their activity when expressed in cell culture and presented with dietary carotenoids. When supplied with the yellow carotenoids zeaxanthin (A, B), beta - carotene (C, D), lutein (E, F), or beta - cryptoxanthin (G, H), CYP2J19 and BDH1L catalyze the production of ketocarotenoid products identified by their retention times (left) and spectra (right) relative to known standards; in these assays pictured here, we also transfected cells with *TTC39B* to enhance ketocarotenoid production for improved visualization of products. However, in none of our assays testing any enzyme and carotenoid combination did we detect carotenoids with the retention times and spectra characteristic of the most abundant ketocarotenoids found in house finch feathers (I, J).

Importantly, our study is the first to test the function of any homolog of these focal genes with the yellow carotenoid β-cryptoxanthin, which has been implicated as a key dietary substrate for red coloration in house finches and the precursor of the major red plumage ketocarotenoid 3-OH-echinenone (Stradi et al. 1996; Hill 2000). If the CYP2J19/BDH1L system mediates red color expression in house finches, then we would expect these enzymes to catalyze the oxidation of β-cryptoxanthin to 3-OH-echinenone. However, cells co-transfected with house finch *CYP2J19* and *BDH1L* and provided the β-cryptoxanthin substrate yielded major products consistent with adonirubin and astaxanthin, but no 3-OH-echinenone (Figures 1, 4, and S5). In fact, no combination of precursor carotenoid substrate and ketocarotenoid-metabolizing enzyme(s) yielded a product with the spectral properties and HPLC retention time that are characteristic of the 3-OH-echinenone that is abundant in red house finch feathers (Figures 1, 4, S3, and S5). This suggests that even if *CYP2J19* and *BDH1L* are expressed in house finch tissues other than the ones we measured, this combination of enzymes does not catalyze the production of 3-OH-echinenone from any of the four most abundant dietary carotenoid precursors observed in house finches (McGraw et al. 2006). Thus, house finches appear to use an as-yet-unknown alternative enzymatic pathway to produce 3-OH-echinenone.

Given this finding, we searched for a new candidate gene that may be involved in carotenoid metabolism in house finches. We cloned and tested nine other genes that were expressed in liver and/or follicle tissue and were enriched in molting males, or had possible links to the types of enzymatic reactions that could produce ketocarotenoids from precursors (Table S3). However, after testing each candidate with at least two of the four main carotenoid substrates found in house finches (β-cryptoxanthin, β-carotene, lutein, and zeaxanthin), we found no evidence of newly metabolized products in any of our assays of candidates (e.g. Figure S6).

### Enzyme Localization

To test the prediction that one or both carotenoid-metabolizing enzymes act within the mitochondria, we generated expression constructs for each protein with a C-terminal florescent tag and expressed these in HEK293 cells along with marker constructs that localize florescent proteins to the mitochondria (Mito-7) or endoplasmic reticulum (ER-3; Olenych et al. 2007). Contrary to our predictions, neither CYP2J19 nor BDH1L co-localized with the mitochondrial marker and were instead widely distributed, appearing to localize to the endomembrane system and partially co-localize with the endoplasmic reticulum marker (Figures 5, S7, and S8). When co-expressed, CYP2J19 and BDH1L strongly co-localized within the cell, suggesting that these two enzymes are found in the same cellular compartment (Figure 5). TTC39B, on the other hand, appears widely distributed throughout the cell in a pattern consistent with cytoplasmic localization (Figure S9). The localization of these heterolgously expressed tagged proteins is consistent with machine-learning-based (MULocDeep) amino acid sequence analysis predictions of a 99.9% probability of ER localization for house finch CYP2J19, 99.4% ER localization for house finch BDH1L, and 77.7% probability of cytoplasmic localization for house finch TTC39B (Jiang et al. 2021; Jiang et al. 2023).

**FIG. 5.**
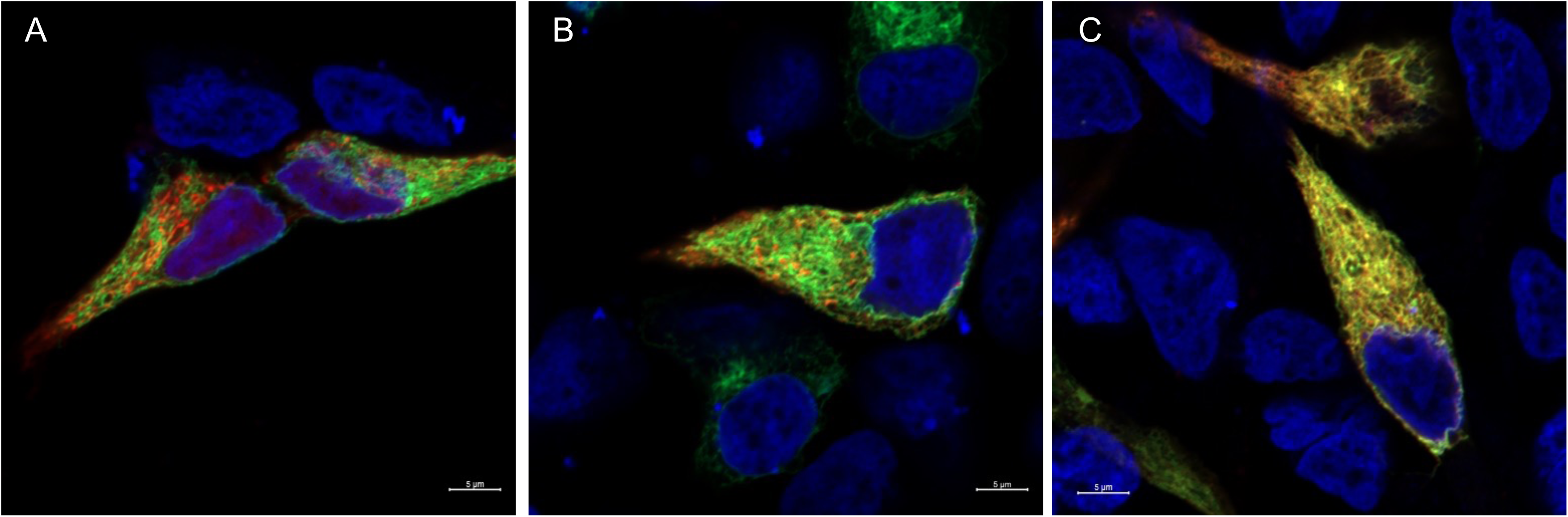
Images of HEK293 cells expressing house finch *CYP2J19* tagged with an in - frame fusion of *GFP*, alongside red fluorescent markers that localize to mitochondrial membranes (A), the endoplasmic reticulum (B), or house finch *BDH1L* with an in - frame fusion of mCherry or *dsRed* (C). Cell nuclei are stained with DAPI and shown in blue. Scale bar = 5 μm.

### Phylogenetic Comparison

Our results suggest that CYP2J19 may not be involved in the expression of red coloration in house finches. Unlike many red bird species that have been previously linked to *CYP2J19*, house finch plumage is primarily pigmented with the asymmetric C4-ketocarotenoids 3-OH-echinenone and 4-oxo-rubixanthin (Figure 2). We conducted carotenoid pigment analysis of red feathers of a purple finch (*Haemorhous purpureus*), which is the sister taxon to the house finch, and observed a similar profile of carotenoids, with 3-OH-echinenone being the predominant ketocarotenoid (Figure S10). To explore how widespread this mechanism of coloration is among all birds, we mapped the occurrence of these asymmetric ketocarotenoids and other major types of carotenoids across bird taxa (Figure 2). Among the species characterized, the co-occurrence of 3-OH-echinenone and 4-oxo-rubixanthin is limited to the red-colored Fringillid finches, like house finches, crossbills (*Loxia* spp.), and rosefinches (*Carpodacus* spp.), while asymmetric ketocarotenoids are found more broadly across *Ramphocelus* tanagers, some *Icterus* orioles, and infrequently among several other songbirds, as well as the Northern flicker (Figure 2). Asymmetric β,β-C4-ketocarotenoids, which include 3-OH-echinenone, echinenone, rupricolin, and adonixanthin, were detected in the feathers of 38/230 (16.5%) of species examined; nearly half of these (17 species) are within the Fringillid finch clade.

## Discussion

We tested key predictions for the enzymes, tissues, and cellular compartments mediating the expression of ketocarotenoid-based ornamental red coloration in house finches. We found that: 1) house finches do not express *CYP2J19* at the putative sites of carotenoid metabolism; 2) in cultured cells, CYP2J19 and BDH1L do not catalyze the formation of the main house finch pigment 3-OH-echinenone as a major product; and, 3) CYP2J19 and BDH1L do not localize to the mitochondria. These observations have important implications for understanding the signal function and evolution of red carotenoid coloration in animals.

When we consider the molecular structures of the dietary carotenoids and the products obtained in our assays of CYP2J19/BDH1L in cultured cells, it becomes clear why birds like house finches that accumulate asymmetric β,β-C4-ketocarotenoids in their plumage likely utilize an alternative enzymatic pathway to CYP2J19/BDH1L. When we provided *CYP2J19-* and *BDH1L*-expressing cells with carotenoid substrates with β-rings at both ends of the molecule (e.g. β-carotene, zeaxanthin, and β-cryptoxanthin), we always observed the symmetric addition of ketone groups at the C4 and C4’ positions. With a lutein substrate that has one β-ring and one ε-ring, there is an asymmetric addition of a ketone group only at the C4 position and the ε-ring is unmodified, yielding α-doradexanthin. In contrast, 3-OH-echinenone has two β-rings, but only one ketone group at the C4 position, and we have never observed this or any other asymmetric β,β-C4-ketocarotenoid product in our assays of the products of CYP2J19/BDH1L. Moreover, house finches appear to express fully functional CYP2J19 and BDH1L in their retina, where yellow dietary carotenoids are metabolized into symmetric ketocarotenoids (like astaxanthin) and used in red cone oil droplets; however, 3-OH-echinenone has never been reported in the avian retina (Goldsmith et al. 1984; Toomey and McGraw 2007; Toomey and McGraw 2009; McGraw and Toomey 2010; McGraw et al. 2013; Toomey et al. 2015; Arteni et al. 2019), further supporting that the CYP2J19/BDH1L does not catalyze the production of asymmetric β,β-C4-ketocarotenoid products.

To our knowledge, the only known instances of enzymes catalyzing the metabolism of symmetric yellow carotenoid substrates into asymmetric β,β-C4-ketocarotenoid products are described in cyanobacteria (Fernández-González et al. 1997; Tsuchiya et al. 2005). In *Synechosystis* sp. and *Gloeobacter violaceus* cyanobacteria, the enzymes CrtO and CrtW, respectively, have been found to catalyze the production of echinenone (an asymmetric β,β-C4-ketocarotenoid) from β-carotene (a symmetric non-ketocarotenoid; Fernández-González *et al*., 1997; Tsuchiya *et al*., 2005). In the house finch, the enzyme pyridine nucleotide-disulfide oxidoreductase domain 2 (PYROXD2) is the most similar to CrtO and CrtW, sharing approximately 35% amino acid identity with *Synechosystis* CrtO. We cloned and tested *PYROXD2* in our cell culture assays, but did not detect any carotenoid metabolizing activity (Figure S6).

The discovery that not all birds with red ketocarotenoid-pigmented plumage utilize the CYP2J19/BDH1L pathway presents an opportunity for new hypotheses to explain diversity in color variability and the potential information communicated by such variation. One of the most remarkable features of carotenoid coloration in male house finches, for example, is the unusually wide variation in hue and carotenoid composition of their ornamental plumage, ranging from dull yellow to bright red (Hill, 2002). There are few bird species in the world with such extreme phenotypic variation in carotenoid coloration. As a comparison to house finches, male northern cardinals (*Cardinalis cardinalis*) have feather pigment profiles dominated by the ketocarotenoid α-doradexanthin, which is consistent with CYP2J19/BDH1L-mediated oxidation of the dietary carotenoid lutein (McGraw et al. 2003; Toomey et al. 2022a). Male cardinals vary in intensity of red, not across a spectrum of yellow to red; even cardinals held on a low-carotenoid diet while molting in captivity grow pale red feathers that contain the same ketocarotenoids as wild birds, but in lower concentrations (McGraw et al., 2001). In contrast, male house finches in captivity on seed diets grow pale yellow feathers lacking any red coloration (Hill 1992). One hypothesis to explain the extreme phenotypic variation of male house finches is that the enzymatic pathway leading to 3-OH-echinenone predisposes these birds to be more variable in ornamental coloration. Indeed, one of the few reported bird species that is as variable as the house finch in carotenoid coloration, the red crossbill (*Loxia curvirostra*), also uses 3-OH-echinenone as its primary red pigment (Stradi et al. 1996) and produces only yellow plumage in captivity (Völker, 1957). These patterns may be related to the fact that 3-OH-echinenone, but not other ornamental ketocarotenoids like α-doradexanthin or astaxanthin, is a vitamin A precursor, potentially leading to flexible use of this ketocarotenoid as a colorant or vitamin A source in species like house finches and red crossbills (Hill and Johnson 2012). Alternatively, the subcellular location or some other attribute of the enzyme pathway to 3-OH-echinenone may predispose birds to have variable expression of ornamental plumage.

The subcellular location of ketocarotenoid metabolism has become a key component of recent hypotheses seeking to explain carotenoid-based color variation. It has been proposed that the enzyme used to produce red ketocarotenoids— assumed to be CYP2J19, after its discovery—localizes to the inner mitochondrial membrane, creating functional links between mitochondrial respiration and the production of red pigments (Johnson and Hill 2013; Hill et al. 2019; Powers and Hill 2021). This hypothesis stemmed largely from the observation that the hepatic mitochondria of house finches contain high concentrations of 3-OH-echinenone (Ge et al. 2015; Hill et al. 2019). However, using tagged proteins, we discovered that not only CYP2J19 but also BDH1L function outside of the mitochondria; the most likely subcellular location of both of these enzymes is the endoplasmic reticulum (ER).

While the ER and mitochondria have clear functional connections (Cohen et al. 2018; Degechisa et al. 2022), the ER in particular plays a major role in lipogenesis and lipid homeostasis (Fu et al. 2012; Stevenson et al. 2016), providing new avenues for exploring the mechanisms driving color variation. Moreover, this observation draws further contrast between house finches, which do not use the CYP2J19/BDH1L pathway, and the birds that likely do, such as the northern cardinal. A preliminary analysis of the hepatic mitochondria of male northern cardinals detected no red coloration, in contrast to what is observed in the mitochondria of house finches (Hill GE and Zhang Y, personal observations). We speculate that the undiscovered enzyme that house finches use to produce ketocarotenoids may localize to the mitochondria, explaining both these observations and the links between 3-OH-echinenone, redness, and mitochondrial function observed in this species (Hill et al. 2019; Koch et al. 2023). Further study of carotenoid localization, subcellular functionality, and gene expression across ketocarotenoid-pigmented taxa will be essential to explore such possibilities and to identify the alternative enzymatic pathway used by birds like house finches.

To better understand the evolutionary context for the alternative pathway used by house finches to produce red ornamental ketocarotenoids, we placed the house finch within a broader phylogeny of avian taxa and mapped the types of metabolized carotenoids that have been reported in plumage. Asymmetric β,β-C4-ketocarotenoids like 3-OH-echinenone have been reported not just in Fringillid finches related to the house finch, but also in the plumage of species belonging to several disparate families of perching birds (Passeriformes), including Icteridae, Thraupidae, and Regulidae, and a very few non-passerine species. This suggests that, like house finches, many avian species employ another mechanism of ketocarotenoid metabolism in addition to or as an alternative to the CYP2J19/BDH1L mechanism. The evolution of multiple mechanisms of metabolism is not necessarily surprising because ketocarotenoid-based coloration is a likely target of sexual selection. Parsing the benefits and costs of alternative or redundant mechanisms and mapping their distribution across the avian phylogeny holds great promise for new directions for understanding avian diversity.

Interestingly, we observed that house finches express *BDH1L* in their growing feather follicles, and in the absence of CYP2J19, BDH1L catalyzes the oxidation of dietary precursor carotenoids into canary xanthophylls. These yellow ornamental carotenoids are often detected at low to moderate concentrations in the carotenoid-pigmented feathers of male house and purple finches, respectively, but are found abundantly in the plumage of yellow Fringillid finches and a wide variety of other avian taxa both with and without red coloration. Our findings are consistent with the possibility that BDH1L plays a role in metabolizing these yellow ornamental pigments, and gains and losses of CYP2J19 and/or alternative enzymes to produce ornamental red ketocarotenoids outside of the retina may explain shifts between yellow and red phenotypes among avian taxa (Ligon et al. 2016; Toomey et al. 2022a; Hooper et al. 2024). Further study will be necessary to test the potential role of BDH1L in the yellow ornamental carotenoids of Fringillid finches and other birds.

In sum, the existence of more than one distinct enzymatic pathway from yellow dietary to red ornamental carotenoid pigments in birds has important implications for understanding honest signaling via carotenoid pigmentation. Hypotheses for the physiological basis of the links between pigmentation and individual quality have shifted in recent years toward an exploration of pathways shared between pigment production and vital cellular processes. With at least two enzyme systems in play, it is no longer reasonable to assume that a single mechanism might accommodate all ketocarotenoid pigmentation across Aves. A better fundamental understanding of the evolution of honest signaling will come from a full characterization of the biochemical processes that give rise to carotenoid ornamentation.

## Acknowledgements

This research was funded by the National Science Foundation (NSF-IOS-2037741, NSF-IOS-2037735, and NSF-IOS-2037739 to GEH, YZ, and MBT, respectively, and NSF-IOS-2224556 to YZ). Ethan Hare (Auburn University) assisted with house finch capture. We thank Russell Ligon for providing the Fringillidae tree. A portion of the computing for this project was performed at the OU Supercomputing Center for Education & Research (OSCER) at the University of Oklahoma (OU).

## Data Accessibility

Differential gene expression statistical results, gene expression data, and sequences of all primers used are available in the supplementary material. Reference sequences for genes tested are available in figshare (doi: 10.6084/m9.figshare.26263130). Transcriptomic raw data are available on NCBI under BioProject PRJNA1076810.

