## Supplemental Figures 1-10 for "Multiple pathways to red carotenoid coloration: House finches (*Haemorhous mexicanus*) do not use CYP2J19 to produce red plumage"

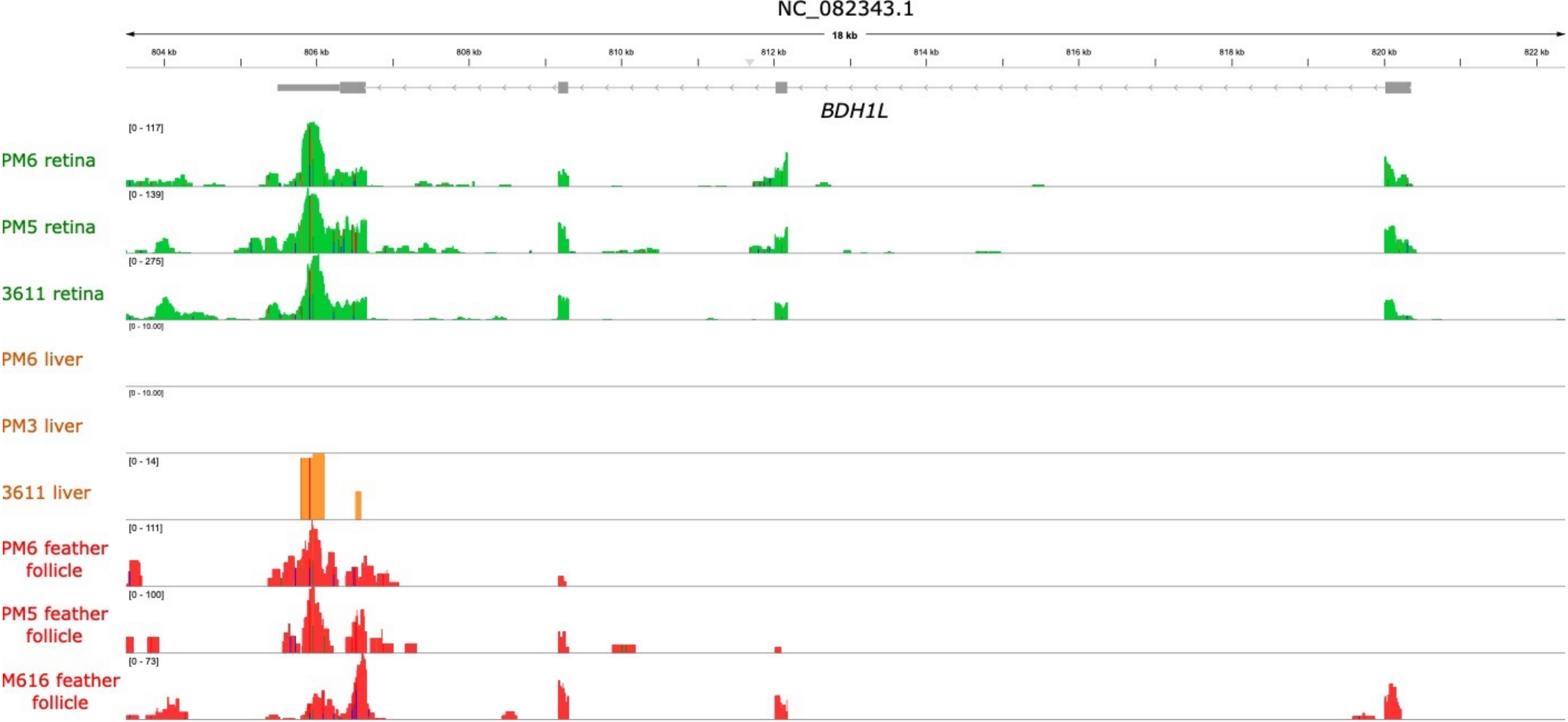


**FIG. S1.** A visualization of expression levels of reads from male house finch retina (top, green), liver (middle, orange), and growing feather follicle (red, bottom) samples for the house finch gene *BDH1L* on chromosome 3. The exons of *BDH1L* are designated in grey boxes.

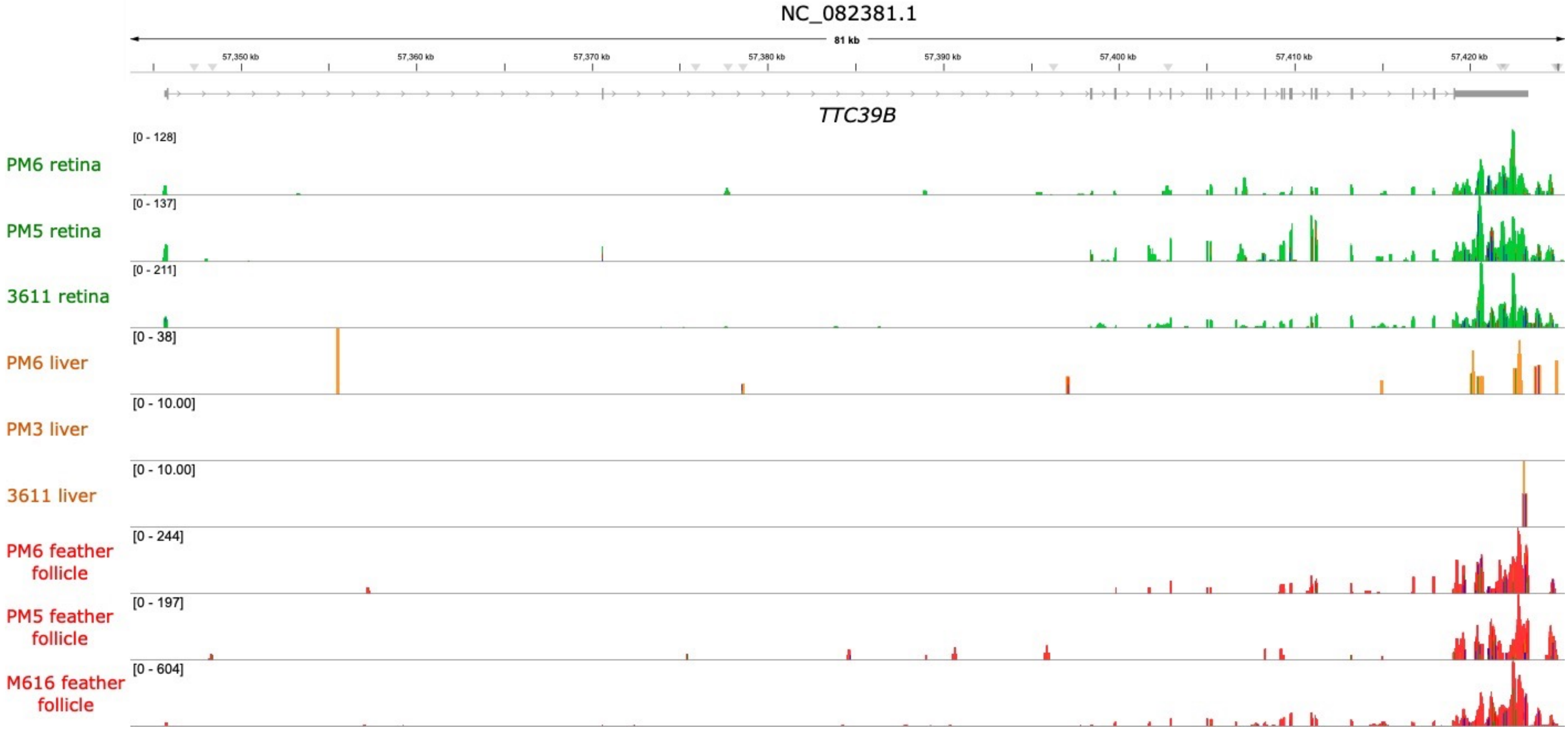


**FIG. S2.** A visualization of expression levels of reads from male house finch retina (top, green), liver (middle, orange), and growing feather follicle (red, bottom) samples for the house finch gene *TTC39B* on the Z chromosome. The exons of *TTC39B* are designated in grey boxes.

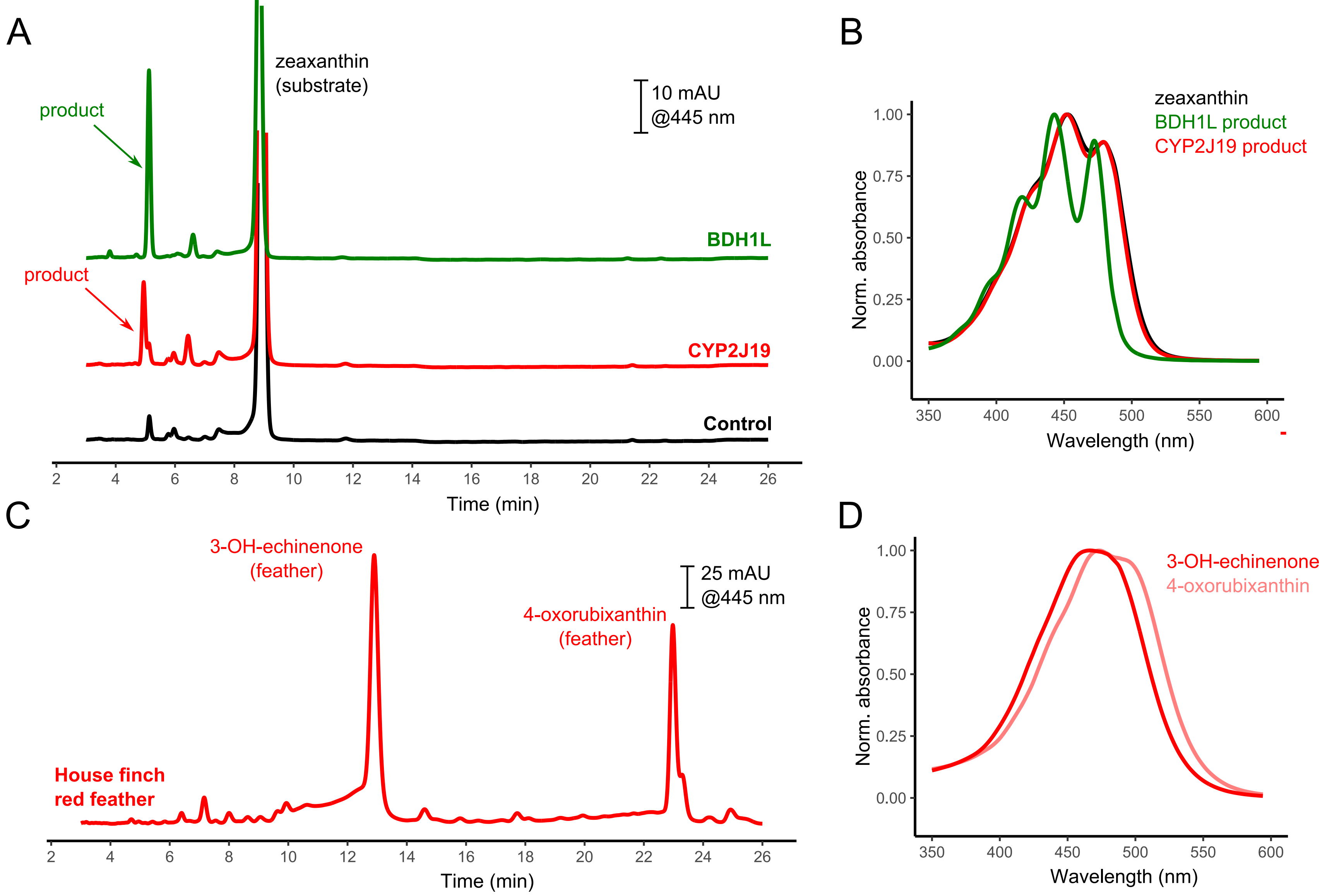


**FIG. S3.** In accordance with previous study of the chicken homologs of these enzymes, house finch CYP2J19 or BDH1L alone catalyze the metabolism of dietary yellow carotenoids (here, zeaxanthin) into modified yellow carotenoids (A, B). The retention times and spectral properties of these single-enzyme products are not consistent with those of the ketocarotenoids present in house finch feathers (C, D).

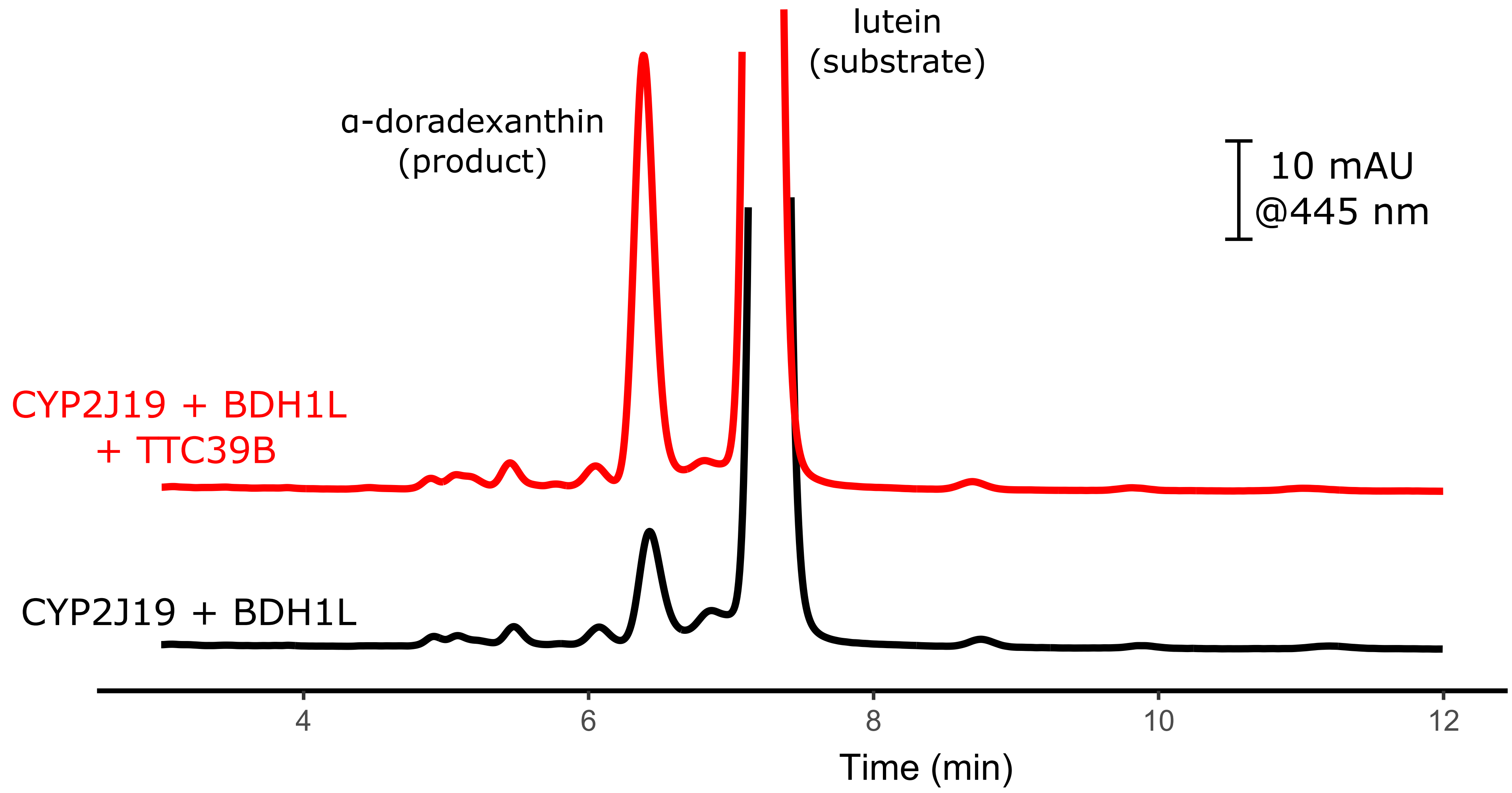


**FIG. S4.** As previously found for the chicken homolog of this gene, the product of house finch *TTC39B* enhances CYP2J19/BDH1L-catalyzed production of ketocarotenoids from dietary carotenoids. Here, cells provided lutein and co-transfected with *CYP2J19* and *BDH1L* produced more of the red ketocarotenoid α-doradexanthin when they were also transfected with *TTC39B*.

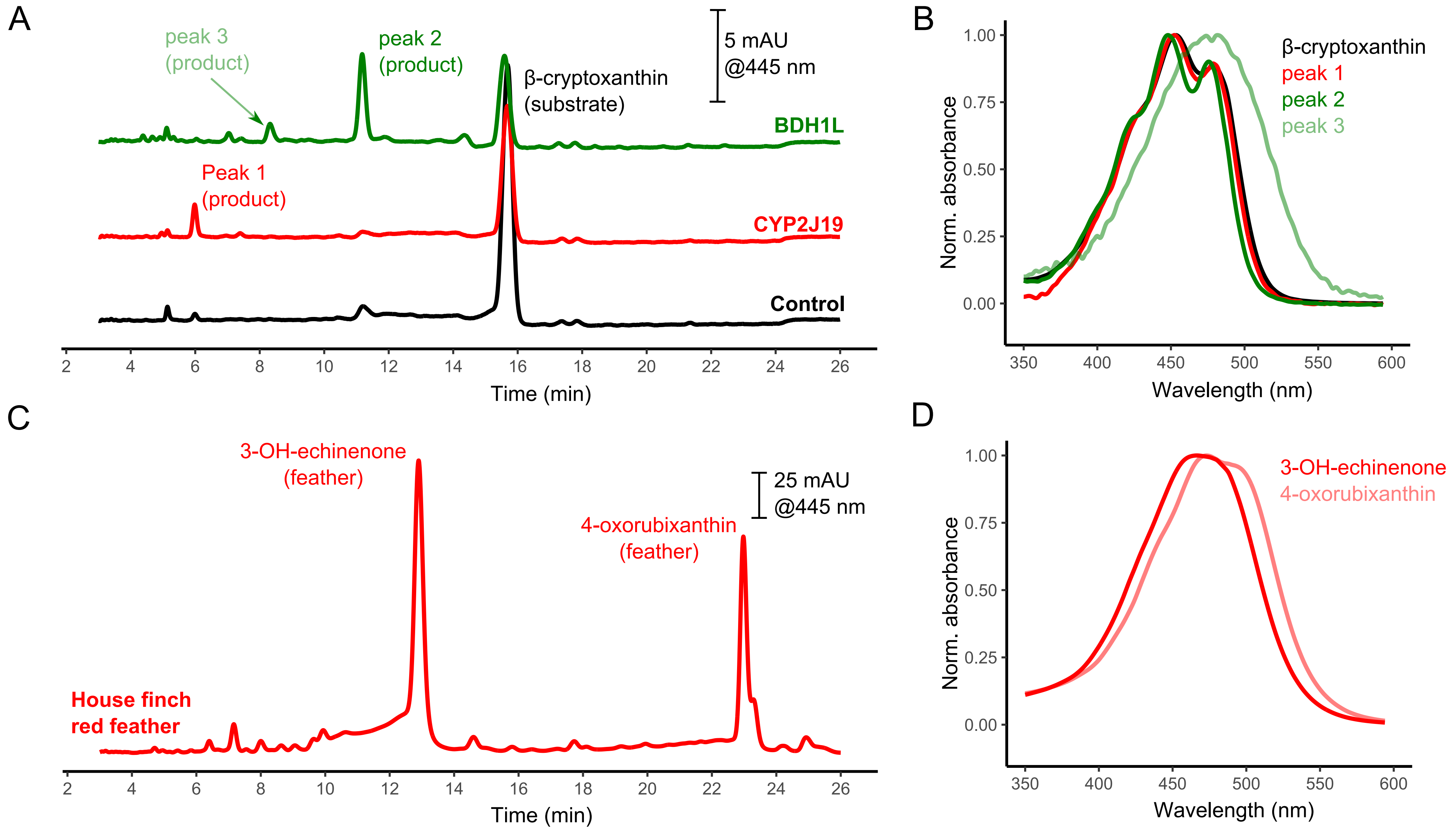


**FIG. S5.** When presented with the dietary carotenoid β-cryptoxanthin, neither BDH1L nor CYP2J19 alone catalyzed the production of a carotenoid (A, B) with properties consistent with the major house finch red ketocarotenoids (C, D). Instead, we detected new products with properties consistent with modified yellow carotenoids.

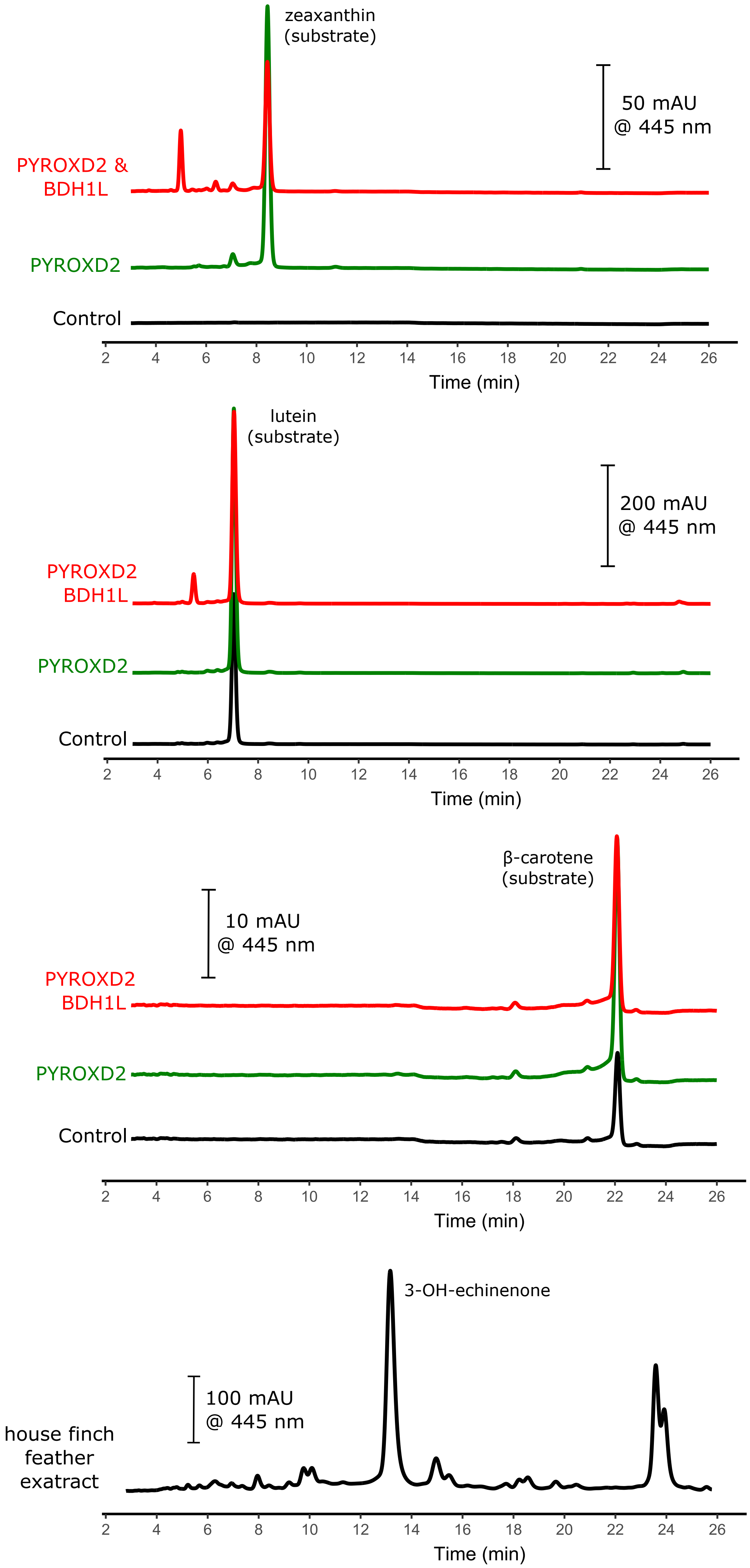


**FIG. S6.** An example of enzymatic activity assay results from one of the candidate genes we tested, *PYROXD2.* We detected no new products when cells expressing *PYROXD2* were presented with any of three different dietary carotenoids; we also co-expressed *PYROXD2* with *BDH1L* in case the candidate enzyme might act on products of BDH1L and lutein or zeaxanthin. In all assays, we transfected cells with *TTC39B* to enhance ketocarotenoid production for ease of visualization, if relevant.

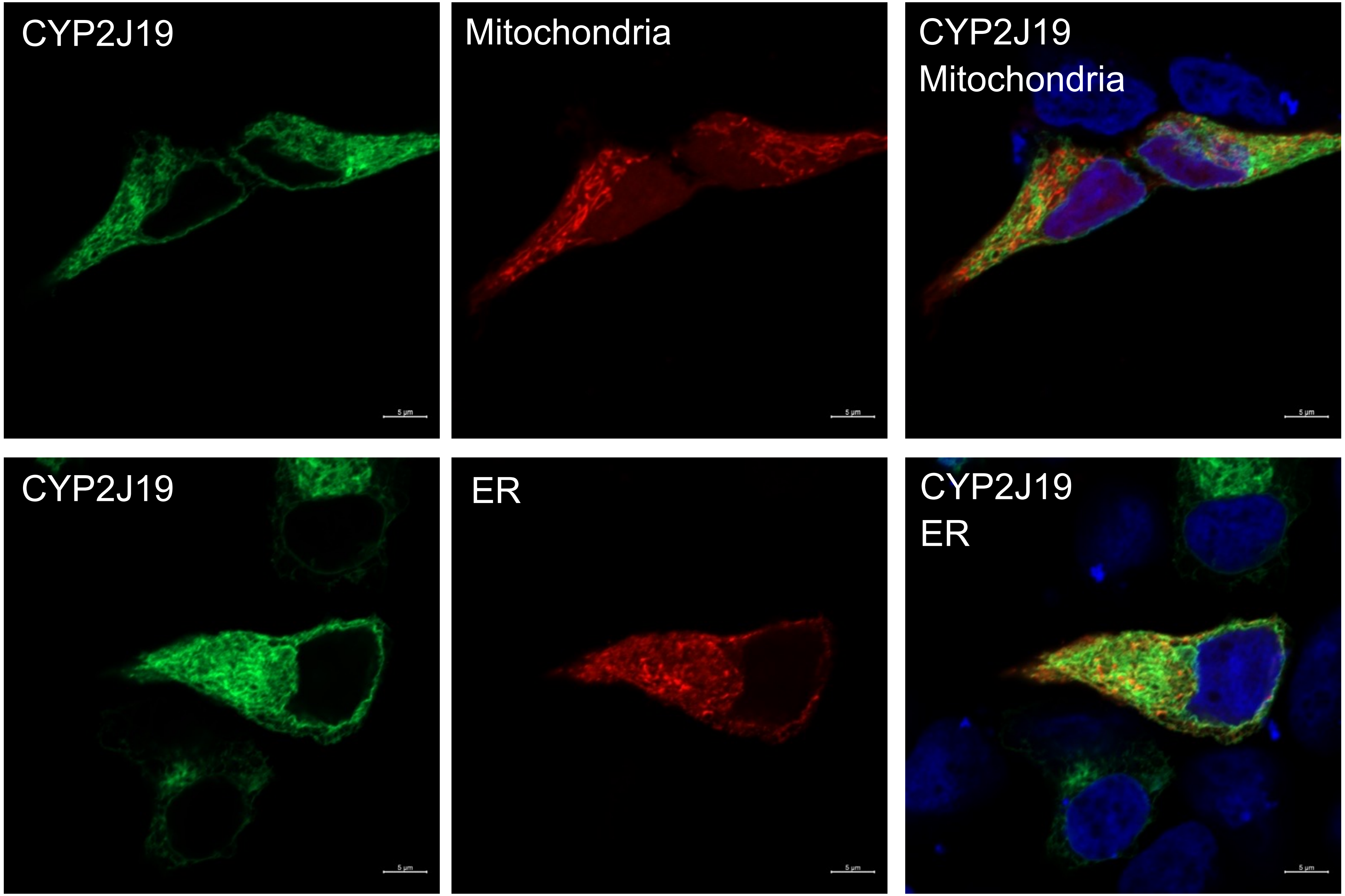


**FIG. S7.** To explore the subcellular localization of CYP2J19, we expressed a construct of this gene with a green fluorescent marker alongside a red fluorescent marker that localizes to mitochondrial membranes (top) or the endoplasmic reticulum (ER; bottom). Cell nuclei are stained with DAPI and shown in blue.

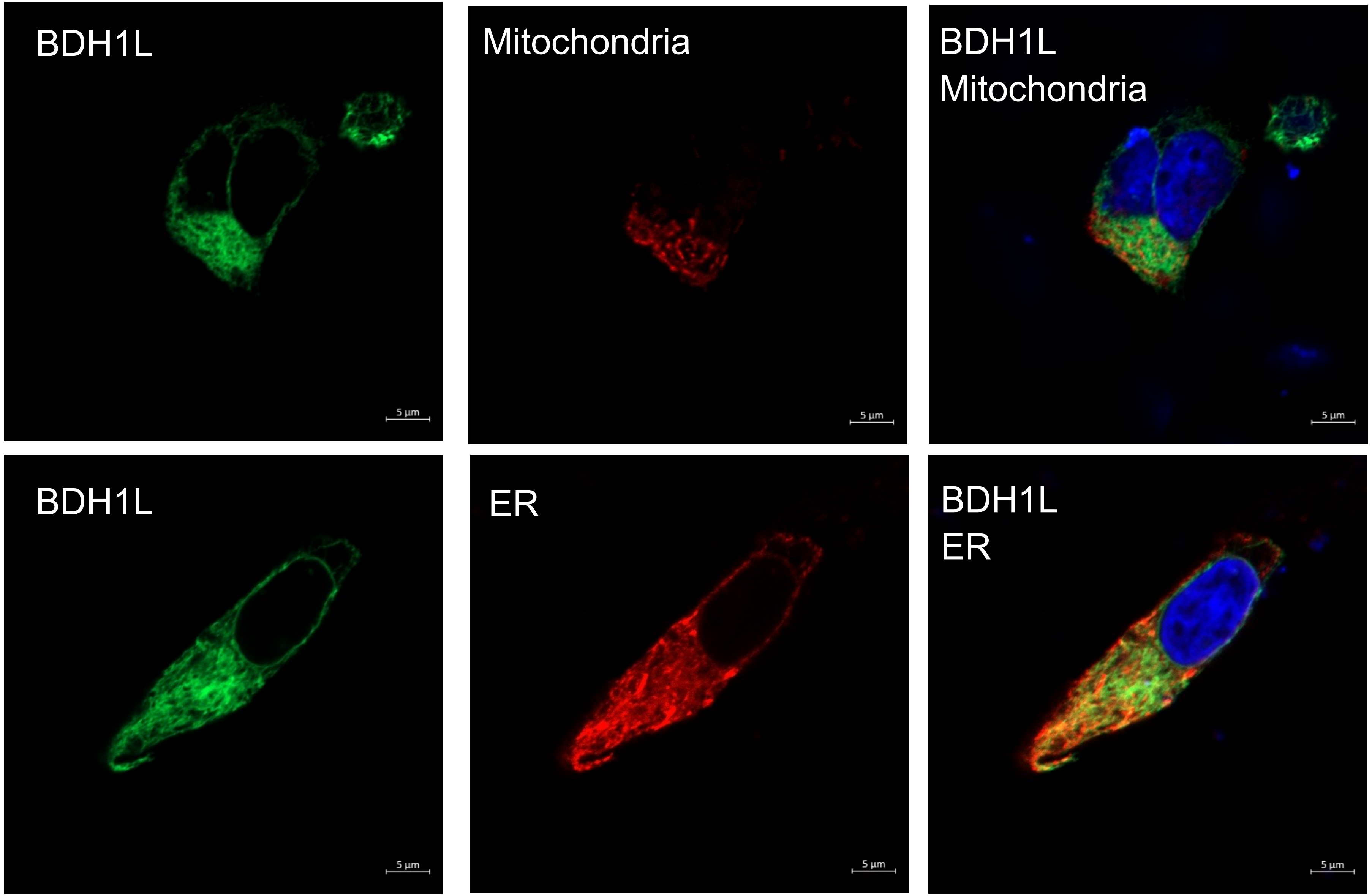


**FIG. S8.** To explore the subcellular localization of BDH1L, we expressed a construct of this gene with a green fluorescent tag alongside a red fluorescent marker that localizes to mitochondrial membranes (top) or the endoplasmic reticulum (ER; bottom). Cell nuclei are stained with DAPI and shown in blue.

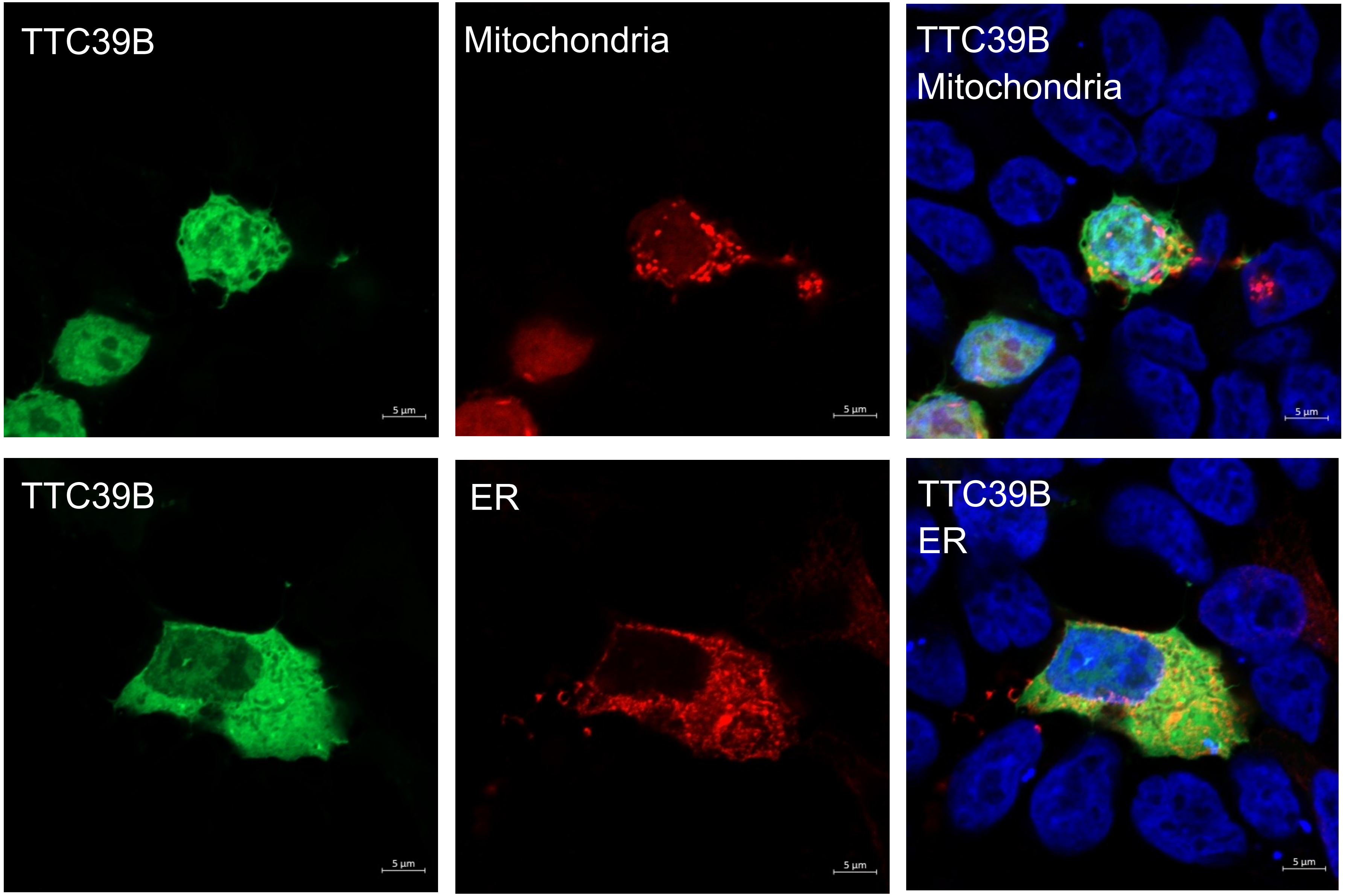


**FIG. S9.** To explore the subcellular localization of TTC39B, we expressed a construct of this gene with a green fluorescent marker alongside a red fluorescent marker that localizes to mitochondrial membranes (top) or the endoplasmic reticulum (ER; bottom). Cell nuclei are stained with DAPI and shown in blue.

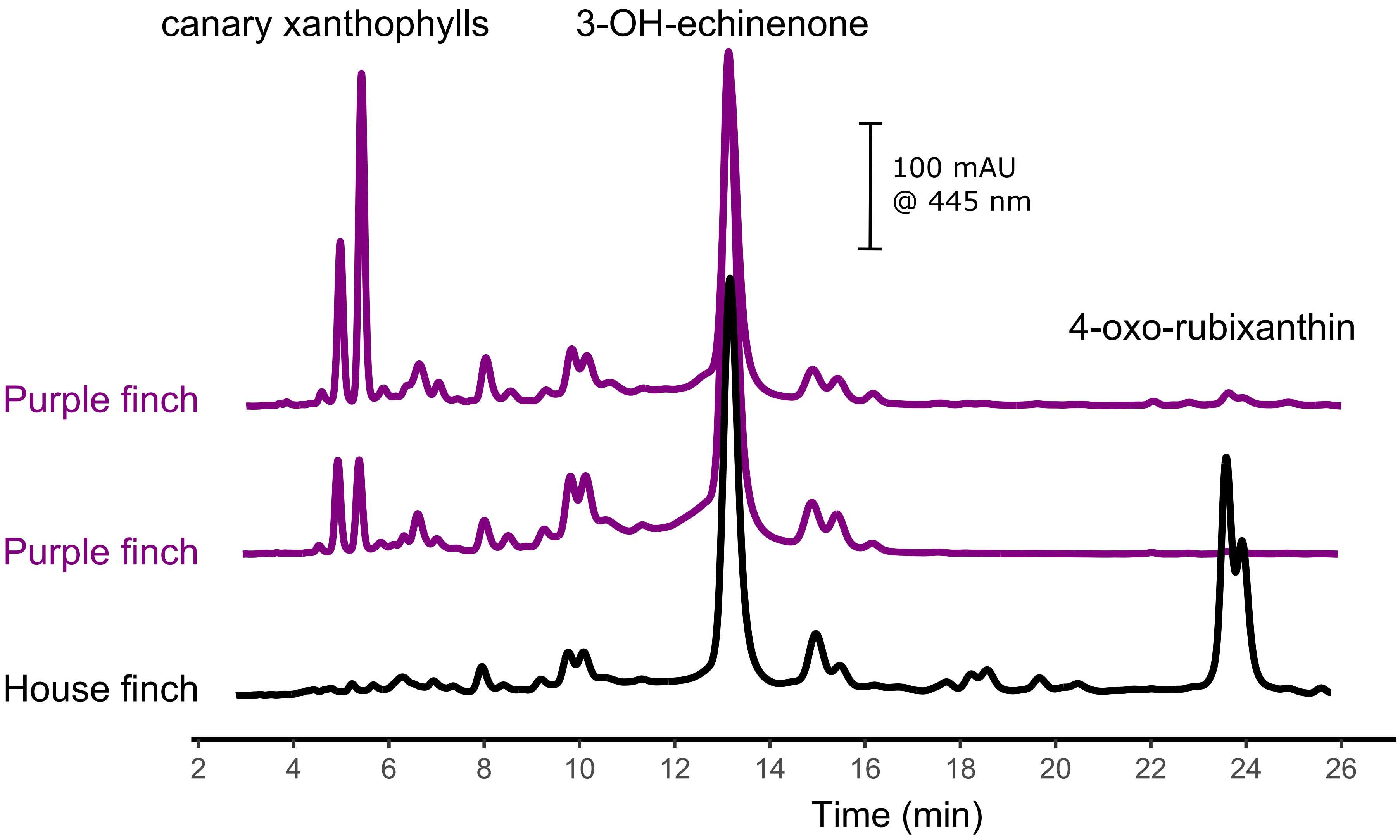


**FIG. S10.** When we analyze carotenoids extracted from feathers of males of the sister species to the house finch, the purple finch *(Haemorhous purpureus*), we find similar profiles as in the house finch. Purple finches, however, appear to deposit greater proportions of canary xanthophylls and less 4-oxo-rubixanthin in their plumage than do house finches.
